# Disinhibition-assisted LTP in the prefrontal-amygdala pathway via suppression of somatostatin-expressing interneurons

**DOI:** 10.1101/799346

**Authors:** Wataru Ito, Brendon Fusco, Alexei Morozov

## Abstract

Natural brain adaptations often involve changes in synaptic strength. The artificial manipulations can help investigate the role of synaptic strength in a specific brain circuit not only in various physiological phenomena like correlated neuronal firing and oscillations but also in behaviors. High and low-frequency stimulation at presynaptic sites has been used widely to induce long-term potentiation (LTP) and depression (LTD), respectively. This approach is effective in many brain areas, but not in the basolateral amygdala (BLA), because the robust local GABAergic tone inside the BLA restricts synaptic plasticity. Here, we identified the subclass of GABAergic neurons that gate LTP in the BLA afferents from the dorsomedial prefrontal cortex (dmPFC). Chemogenetic suppression of somatostatin-positive interneurons (Sst-INs) enabled the *ex vivo* LTP by high-frequency stimulation of the afferent, but the suppression of parvalbumin-positive interneurons (PV-INs) did not. Moreover, optogenetic suppression of Sst-INs with Arch also enabled LTP of the dmPFC-BLA synapses both *ex vivo* and *in vivo*. These findings reveal that Sst-INs but not PV-INs gate LTP in the dmPFC-BLA pathway and provide a method for artificial synaptic facilitation in BLA.

## 1 Introduction

Circuit interrogation using the optogenetics and chemogenetics has become a standard approach for testing the causal role of specific neuronal populations and synapses in brain activities and animal behaviors. The techniques employ depolarizing or hyperpolarizing neuronal compartments, like the soma, dendrites, and synaptic terminals, to trigger or suppress the action potentials and release of neurotransmitter (1, 2). Meanwhile, the natural neuronal adaptations driven by experience and learning, or observed during development or in disease, involve brain alterations, not only in the neuronal activity but also in the synaptic efficacy. For example, the auditory fear conditioning, a model of adaptive defensive behavior, strengthens the auditory inputs to the lateral amygdala (3), whereas the fear extinction training depresses the prefrontal-amygdala synapses and strengthens the reciprocal amygdala-prefrontal synapses (4, 5). Modeling and quantitative analyses of such naturally occurring brain adaptations require methods for selective manipulation of the synaptic strength both *ex vivo* and *in vivo*.

By applying high- or low-frequency presynaptic stimulation, synapses, in many cases, can be potentiated or depressed, respectively (6). This simple technique has been employed in optogenetics for manipulating synaptic strength *ex vivo* and *in vivo* and proved successful in several cases. LTP was obtained in the recurrent synapses in hippocampal area CA3 by applying 20 Hz stimulation (7) and in the cortico-striatal synapses by applying the theta-burst stimulation (8). LTD was obtained in the inputs to the nucleus accumbens from the infralimbic cortex and basolateral amygdala by applying the low-frequency presynaptic stimulation (9, 10). At the same time, some synapses do not follow the frequency rule—the high-frequency stimulation generated LTD in the inputs from BLA to the dorsomedial prefrontal cortex (dmPFC) (11).

The synaptic strength in the BLA afferents has been recognized as a critical determinant of fear behaviors and readily changes by emotional experiences. For example, the auditory fear conditioning, in which the animal experiences a neutral conditioned stimulus (CS) followed by electrical footshocks as the unconditioned stimulus (US), facilitates the remote inputs to the lateral subdivision of BLA (3, 12, 13). However, an artificial LTP induction in the BLA afferents, by solely presynaptic stimulation, was predicted to be ineffective because the local GABAergic neurons provide potent feedforward inhibition and gate plasticity in the remote glutamatergic inputs (14, 15). Therefore, the robustness of the “natural” behavior-driven plasticity is explained by the US actions to disinhibit the BLA by attenuating GABAergic transmission via multiple mechanisms, including the secretion of neuromodulators (15-17).

Here, to achieve reliable artificial LTP inductions in the BLA input from dmPFC, which is the critical circuit for emotional learning and control of mood, we tested the effects of chemogenetic/optogenetic suppression of GABAergic transmission during the LTP induction by high-frequency stimulation. The two major classes of GABAergic neurons—the parvalbumin-positive (PV-INs) and somatostatin-positive interneurons (Sst-INs) (18)—were suppressed individually, which revealed that the Sst-INs gate the artificially-induced LTP.

## 2 Materials and methods

### 2.1 Animals

All mice were either wildtype or transgenic males on the 129SvEv/C57BL/6N F1 hybrid background. To obtain the mice expressing hM4Di (19) in Sst-INs or PV-INs, homozygous R26-LSL-Gi-DREADD males (JAX Stock No: 026219) on C57BL/6N background were crossed with homozygous interneuron-specific Cre driver females on 129SvEv background—either the Sst-IN-specific Cre driver (Sst-Cre), *Sst*^*tm2.1(cre)Zjh*^ (20) (JAX: 013044) or the PV-IN-specific Cre driver (PV-Cre), *Pvalb*^*tm1(cre)Arbr*^ (21) (JAX: 008069). To obtain heterozygous Sst-Cre mice, wild type C57BL/6N males were crossed with the 129SvEv homozygous *Sst*^*tm2.1(cre)Zjh*^ females. All breedings were the trios of one male and two females on the C57BL/6N and 129SvEv backgrounds, respectively. Male pups were weaned at p21-p25 and housed 3-5 littermates per cage. All experiments were approved by Virginia Tech IACUC and followed the NIH Guide for the Care and Use of Laboratory Animals.

### 2.2 Surgery—Viral Injection and Optrode Implantation

AAVs for expressing Chronos or Cre-activated Arch were generated from pAAV-Syn-Chronos-GFP (22) (Addgene #59170) or pAAV-FLEX-Arch-GFP (Addgene #22222), respectively, gifts from Edward Boyden. The viruses (pseudotype 5 for Chronos and pseudotype 1 for Arch) were prepared by the University of North Carolina Vector Core (Chapel Hill, NC). At p28, the heterozygous Sst-Cre male mice were anesthetized by intramuscular injection of Ketamine/Xylazine/Acepromazine, 100/5.4/1 mg/kg, placed in a stereotaxic apparatus (David Kopf, Tujunga, CA), and underwent minimum craniotomy (∼0.5 mm diameters). For the dmPFC virus injection, the dura mater was preserved. A heater-pulled short-taper glass pipette (shaft: 0.6/0.4 mm external/internal diameter, beveled tip: 50 µm, diameter, Drummond, Broomall, PA) filled with the virus solution (10^12^ viral particles/ml) was slowly lowered to the target (1.3 mm anterior, 0.4 mm lateral from bregma, and 1.3 mm ventral from brain surface). 0.5 µL of the solution were injected bilaterally at the rate of 0.2 µL/min using a syringe pump connected to the pipette through plastic tubing filled with water as described (23). For the BLA virus injections, the dura mater was removed to allow straight penetration by a less rigid long-taper pipette. 0.4 µL of the virus solution (10^12^ particles/ml) was injected bilaterally at the rate of 0.1 µL/min at the coordinates (1.2 mm posterior, 3.2 mm lateral from bregma, and 4.2 mm ventral from brain surface). The optrodes for *in vivo* recording were fabricated and implanted in BLA at p60 as described (24). For post-operation analgesia, ketoprofen (5 mg/kg) was administered subcutaneously.

### 2.3 Ex Vivo Recordings

#### 2.3.1 General

Mice were anesthetized with intraperitoneal injection of Avertine, 0.4 mg/kg, and intracardially perfused with ice-cold partial sucrose artificial cerebrospinal fluid (ACSF) solution containing (in mM) 80 NaCl, 3.5 KCl, 4.5 MgSO_4_, 0.5 CaCl_2_, 1.25 H_2_PO_4_, 25 NaHCO_3_, 10 glucose, and 90 sucrose equilibrated with 95 % O_2_ / 5 % CO_2_ (25). Amygdala slices, 300 µm thick, were prepared and stored as described earlier (24). Recording chamber was superfused at 2 ml/min with ACSF equilibrated with 95 % O_2_ / 5 % CO_2_ and containing (in mM) 119 NaCl, 2.5 KCl, 1 MgSO_4_, 2.5 CaCl_2_, 1.25 H_2_PO_4_, 26 NaHCO_3_, and 10 glucose (pH 7.4), and maintained at 30 ± 1 °C. Whole-cell recordings were obtained with EPC-10 amplifier and Pulse v8.76 software (HEKA Elektronik, Lambrecht/Pfalz, Germany). Putative glutamatergic neurons in BLA were identified by their pyramidal morphology (26) under Dodt gradient contrast optics (custom made) at 850 nm LED illumination (Thorlabs, Newton, NJ). GABAergic neurons expressing hM4Di-Citrine or Arch-GFP were identified by fluorescence. The recording pipettes (3-5 MΩ) were filled with (in mM) 120 K-gluconate, 5 NaCl, 1 MgCl_2_, 10 HEPES, 0.2 EGTA, 2 ATP-Mg and 0.1 GTP-Na for current-clamp recordings or with 120 Cs-methanesulfonate, 5 NaCl, 1 MgCl_2_, 10 HEPES, 0.2 EGTA, 2 ATP-Mg, 0.1 GTP-Na, and 10 mM QX314 for voltage-clamp recordings. Both internal solutions were set at pH 7.3 and osmolarity 285 Osm. Membrane potentials were corrected by the junction potential of 12 mV. Series resistance (Rs) was 10 – 20 MΩ and monitored throughout experiments to exclude the recording data if the Rs changed more than 20%. LFP recordings were obtained using Multiclamp 700B amplifier and Digidata 1440A (Molecular Device, Sunnyvale, CA). The recording pipettes (1-2 MΩ) were filled with ASCF. Light pulses of 470 nm and 560 nm were generated using LED lamps (Thorlabs) and custom LED drivers based on MOSFET and were delivered through a 40× objective lens (Olympus, Center Valley, PA) at the irradiance of 0.5 to 5 mW/mm^2^, calibrated by a photodiode power sensor (Thorlabs) at the tip of the lens.

#### 2.3.2 LTP

In both the whole-cell and LFP recording, the strength of test pulses (1 ms duration) was adjusted to elicit responses at 30-40% of the maximum. In the whole-cell recordings, test pulses were given every 30 s. LTP was induced by six 2 sec trains of 50 Hz 1 ms pulses. The trains were given at the 10 s interval (Fig.2A). In the LFP recordings, test pulses were given every 20 s. LTP was induced using the “spaced protocol.” It included pairs of 1-sec trains of 50 Hz 1 ms pulses, separated by 10 s. The pairs were repeated five times at the 3 min interval (Fig.3A). This protocol is the same as in a published study on LTP in BLA (27), except the stimulation frequency was decreased from 100 to 50 Hz to allow reliable activation of Chronos (22). In some experiments, continuous yellow light was given during the trains of the blue light pulses. The yellow light strength was set below the levels that trigger the release of glutamate from the axonal terminals expressing Chronos (data not shown).

**Fig.1.**
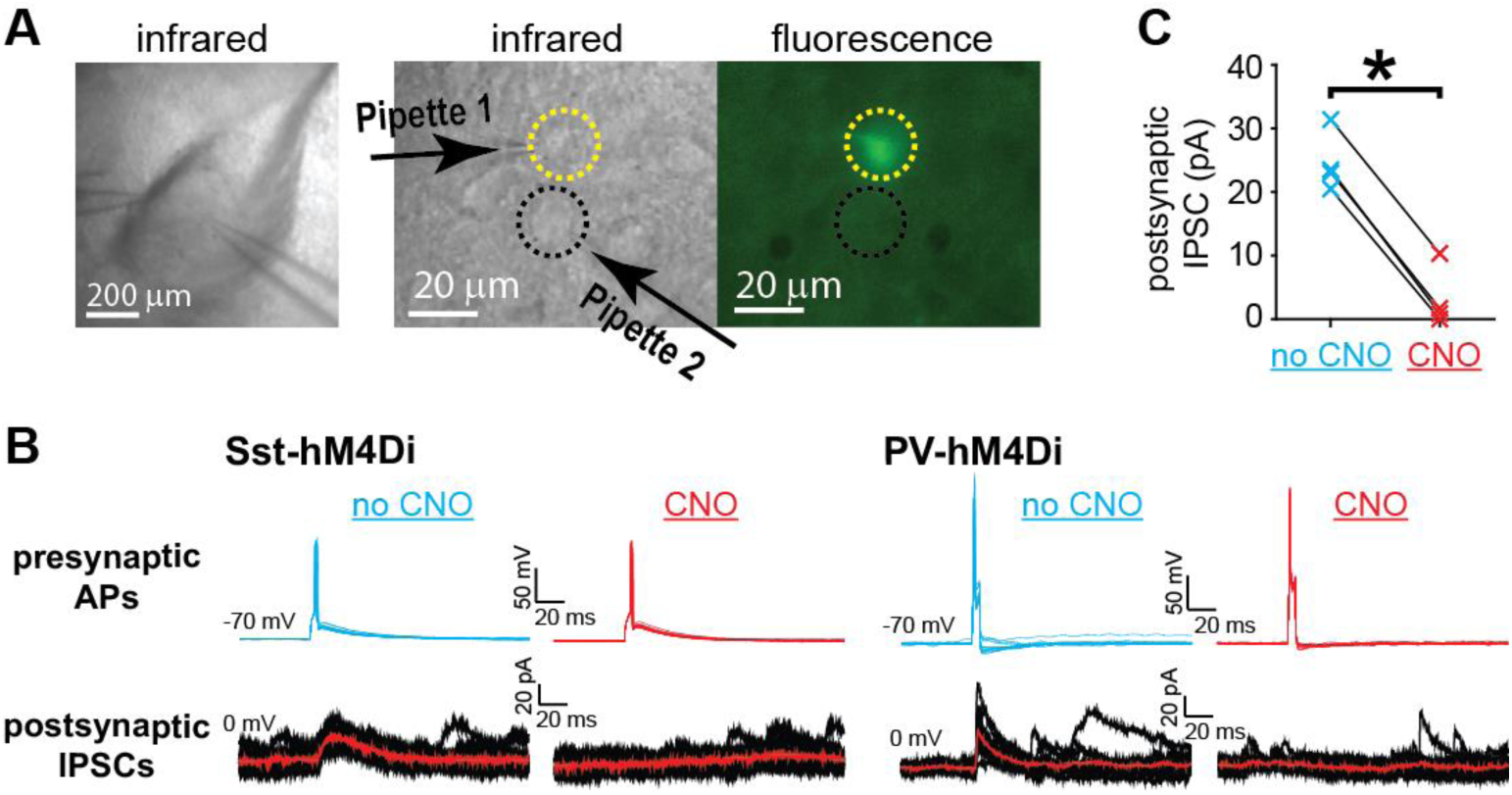
DREADD suppresses GABA release from BLA interneurons during action potentials. (A) Example of the paired whole-cell recording from a BLA slice. Left: Infrared (IR) image at low magnification. Right: high magnification IR and fluorescent images. Dotted yellow and black circles indicate an Sst-IN expressing hM4D(Gi)-citrine identified by fluorescence and a putative principal neuron (PN), respectively. (B) Examples of double patch recordings of APs evoked by current injection in the interneurons (400 pA, every 15 s) (upper) and of the corresponding IPSCs in the connected PNs (lower), in the absence of CNO (no CNO, blue) and after 10 min perfusion with 1 μM CNO (CNO, red). Fifteen traces and IPSC averages (red line) are shown for a pair with an Sst-IN (left, Sst-hM4(Di)) and a pair with a PV-IN (right, PV-hM4(Di)). (C) Summary data for IPSC amplitudes (n=4, including three pairs with Sst-INs and one pair with PV-IN) in the absence and presence of CNO. *p<0.05, Mann Whitney test.

**Fig.2.**
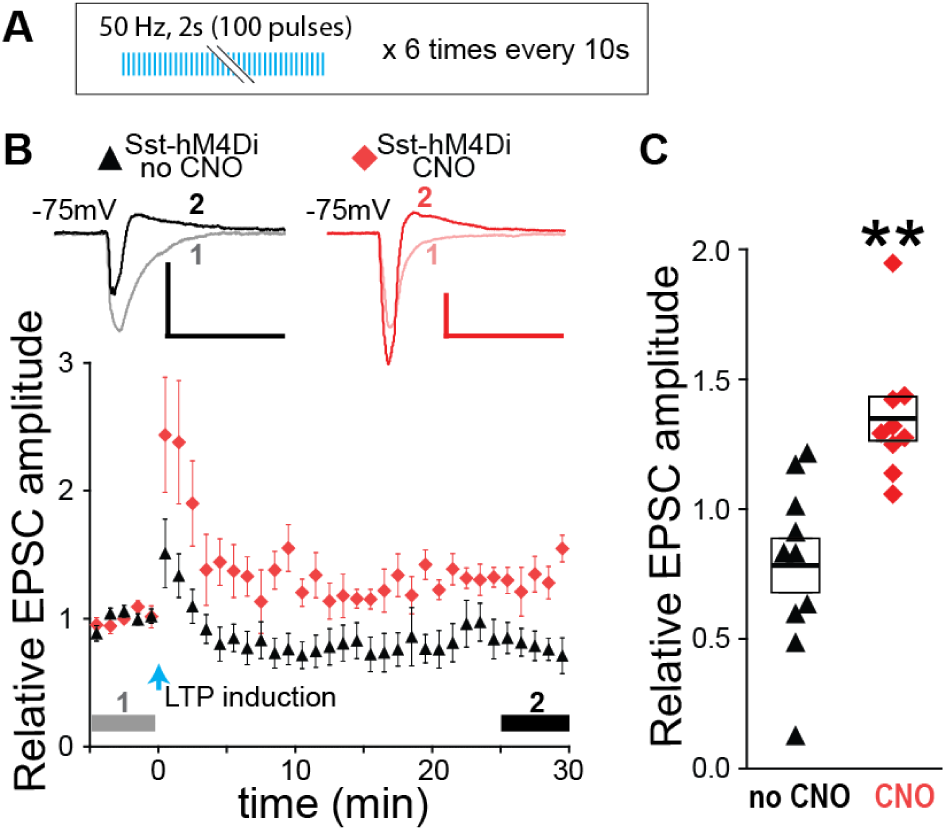
DREADD suppression of Sst-INs enables facilitation of EPSCs in dmPFC-BLA pathway. (A) LTP induction protocol. (B) Relative EPSC amplitudes. Symbols (black triangles: no CNO, red diamonds: CNO) represent the average amplitudes of 2 consecutive EPSCs recorded during each minute. Upper insets: examples of averaged EPSCs before (1) and after (2) LTP induction as indicated by horizontal grey and black bars, respectively. Scales: 100 pA, 50 ms. (C) Relative EPSC amplitudes averaged during (2) for each neuron. n=10 (no CNO) and 9 (CNO). **p<0.01, compared to baseline, one-sample t-test. Boxes and the thick bars inside represent SEM and means.

**Fig.3.**
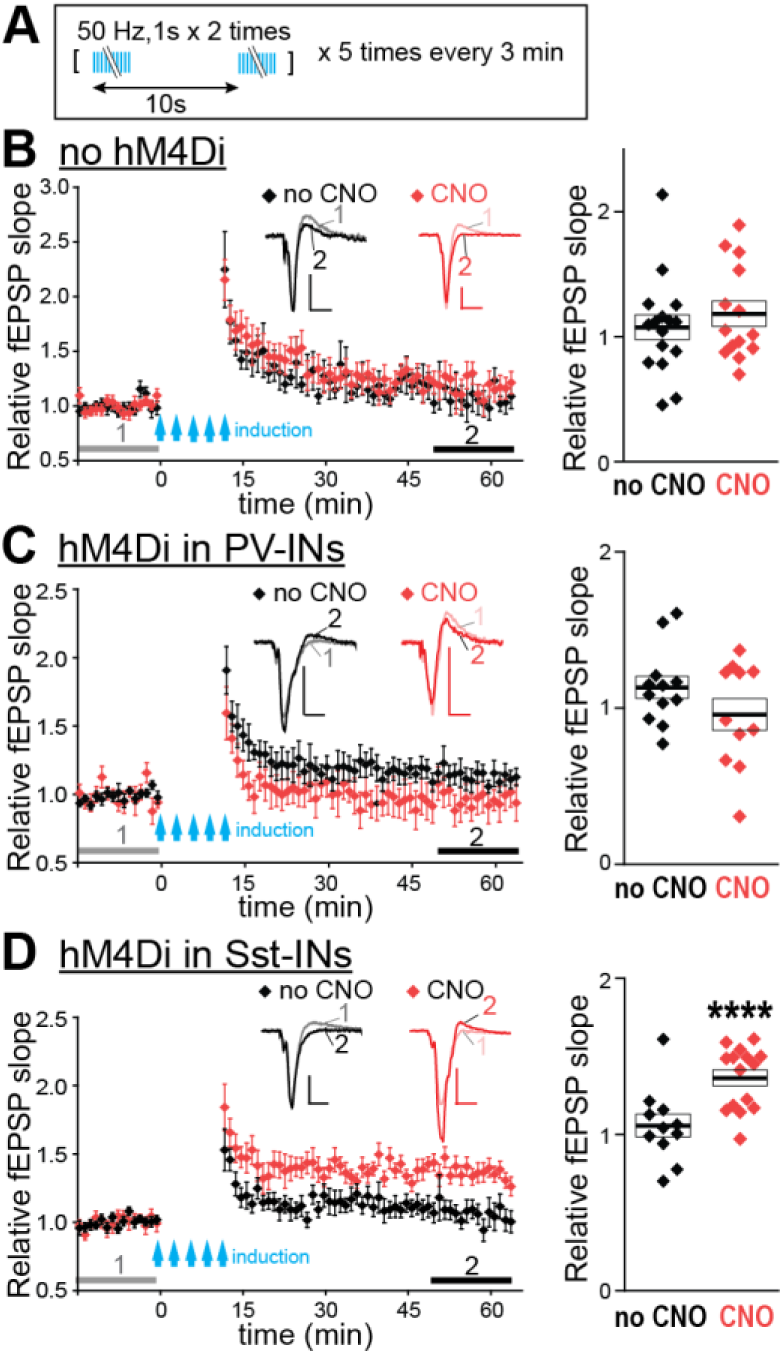
DREADD suppression of Sst-INs enables facilitation of LFPs evoked in the dmPFC-BLA pathway. (A) LTP induction protocol. (B-D) LTP experiments on slices without hM4Di (B), with hM4Di expressed in PV-INs (C), and with hM4Di expressed in Sst-INs (D). Left: Relative fEPSP slopes. Symbols on the diagram represent the averages of three consecutive data points obtained every 20 sec. Insets represent examples of averaged fEPSPs before (1) and after (2) LTP induction as indicated by horizontal grey and black bars. Scales: 0.1 mV, 10 ms. Right: Relative fEPSP slopes averaged during (2) for each slice. n=16 (no CNO) and 14 (CNO) in (B). n=12 (no CNO) and 11 (CNO) in (C). n=11 (no CNO) and 15 (CNO) in (D). ****p<0.0001, compared to baseline, one-sample t-test. Boxes and the thick bars inside represent SEM and means.

### 2.4 In Vivo Recordings

The subject animals, bilaterally injected with the AAV-Chronos in the dmPFC, AAV-Arch in BLA, and bilaterally implanted with the optrodes in the BLA, were housed with the littermates until the experiment. Using the RHA2000-Series Amplifier USB Evaluation Board (RHA2000-EVAL, Intan Technologies), the local field potentials (LFPs) were recorded from BLA of the subject mouse in the home cage without the lid, from where the cagemates were removed temporarily for the duration of the recording. Mice were habituated to the recording environment by connecting to the recording system for 2-3 h per day during 2-3 consecutive days. fEPSPs were elicited in BLA by blue light stimulation of dmPFC terminals expressing Chronos. The strength of the test pulses (1 ms, 2-3 mW at the tip of optrode) was adjusted to obtain the fEPSP slope at 30-40% of the maximum. The LED driver (PlexBright LD-1, Plexon) was analog-modulated by DAQ (Analog Shield, Digilent). The LED driving current was routed to the optrodes in either hemisphere by electrical relays (Arduino 4 relays shield, Arduino). Arduino with a custom Arduino sketch controlled both the DAQ and the relays to give the light stimulation on each side every 30 s, alternating the sides every 15 s. Once the baseline of evoked fEPSPs stabilized, LTP was induced by the same blue light stimulation protocol as in the LFP LTP experiments *ex vivo* (Fig.3A), except that the protocol was repeated three times every one hour. The positions of the optrodes were confirmed by histological analysis.

### 2.5 Data Analysis

Data were processed using custom scripts written in MATLAB (MathWorks) and Clampfit software (Molecular Devices). Statistical analyses were performed using GraphPad Prism 5 (GraphPad Software, La Jolla, CA). Normality was tested using the Shapiro-Wilk test. Datasets with normal distribution were compared using the one-sample t-test. The datasets with non-normal distribution were analyzed using the Mann-Whitney test and the Wilcoxon Signed Rank Test. The difference was deemed significant with p<0.05.

## 3 ‘Results

### 3.1 DREADD-hM4D(Gi) Suppresses GABA Release from BLA Interneurons

The efficiency of DREADD suppression was tested by double-patch recording from connected pairs of an interneuron (IN) expressing hM4Di-Citrine identified by the fluorescence and a putative principal neuron (PN). The brief depolarizing current was injected in the IN to trigger single action potential (AP). It resulted in the inhibitory postsynaptic current (IPSC) in the connected PN. Including CNO (1 µM) in the bath did not prevent APs but diminished IPSCs (Fig.1). The DREADD suppression of presynaptic GABA release despite the presence of action potential was consistent with published findings (28).

### 3.2 Chemogenetic Suppression of Sst-INs Enables LTP Induction Ex Vivo

For faithful activation of dmPFC axons at high-frequency, a fast opsin Chronos (22) was expressed in dmPFC. The 50 Hz trains of light pulses were used for LTP induction (Fig.2A). First, we examined the effect of DREADD suppression of Sst-INs on LTP, by whole-cell recording from PNs in BLA slices expressing hM4Di in Sst-INs. In the absence of CNO, the 50 Hz stimulation of dmPFC axons caused a brief post-tetanic potentiation of the excitatory postsynaptic currents (EPSCs), followed by a rapid EPSC decline with a tendency towards depression at the minutes 25-30 after the induction (p=0.068). In the presence of CNO, the stimulation caused increases in EPSCs lasting for at least 30 min (Fig.2), suggesting that suppression of Sst-INs enables LTP induction.

Repeated neuronal stimulation over extended time intervals, or the spaced LTP protocols, induces LTP more effectively than the shorter protocols, which is consistent with greater efficiency of spaced over massed training (29). However, our attempts to induce LTP with a spaced protocol during the whole-cell recording have failed (data not shown), presumably because the dialysis of the intracellular content limits the time between obtaining the whole-cell configuration and effective LTP induction. To overcome this limitation, in the following experiments, LTP was tested by recording local field potentials (LFPs) and using the spaced LTP protocol (Fig.3A). We run the CNO control, and then tested the effects of suppressing PV-INs, and re-examined the effect of suppressing Sst-INs.

For CNO control, we recorded from slices that did not express hM4Di. There was no significant LTP in the absence or presence of CNO, but there was a tendency towards LTP with CNO (p=0.09) (Fig.3B). In slices with hM4Di in PV-INs, there was no significant LTP in the absence or presence of CNO but a tendency towards LTP in the absence of CNO (p=0.09) (Fig.3C). In slices expressing hM4Di in Sst-INs, there was a significant LTP in the presence of CNO and no LTP in the absence of CNO (Fig.3D). These data indicate that a) CNO in the absence of hM4Di has a minor effect if any on LTP induction, b) suppression of PV-INs does not aid LTP induction, but may rather impede it, and c) suppression of Sst-INs enables LTP.

### 3.3 Arch in Sst-INs Enables LTP Induction Ex Vivo and *In Vivo*

To examine the effects of optogenetic suppression of Sst-INs, Arch was expressed in the Sst-neurons of BLA (Fig.4A). Presynaptic stimulation was given through Chronos expressed in dmPFC terminals in the same way as in the DREADD suppression experiments. Paired recordings confirmed that Arch attenuated GABAergic transmission between Sst-IN and principal neuron (PN) in BLA (Fig.S1).

**Fig.4.**
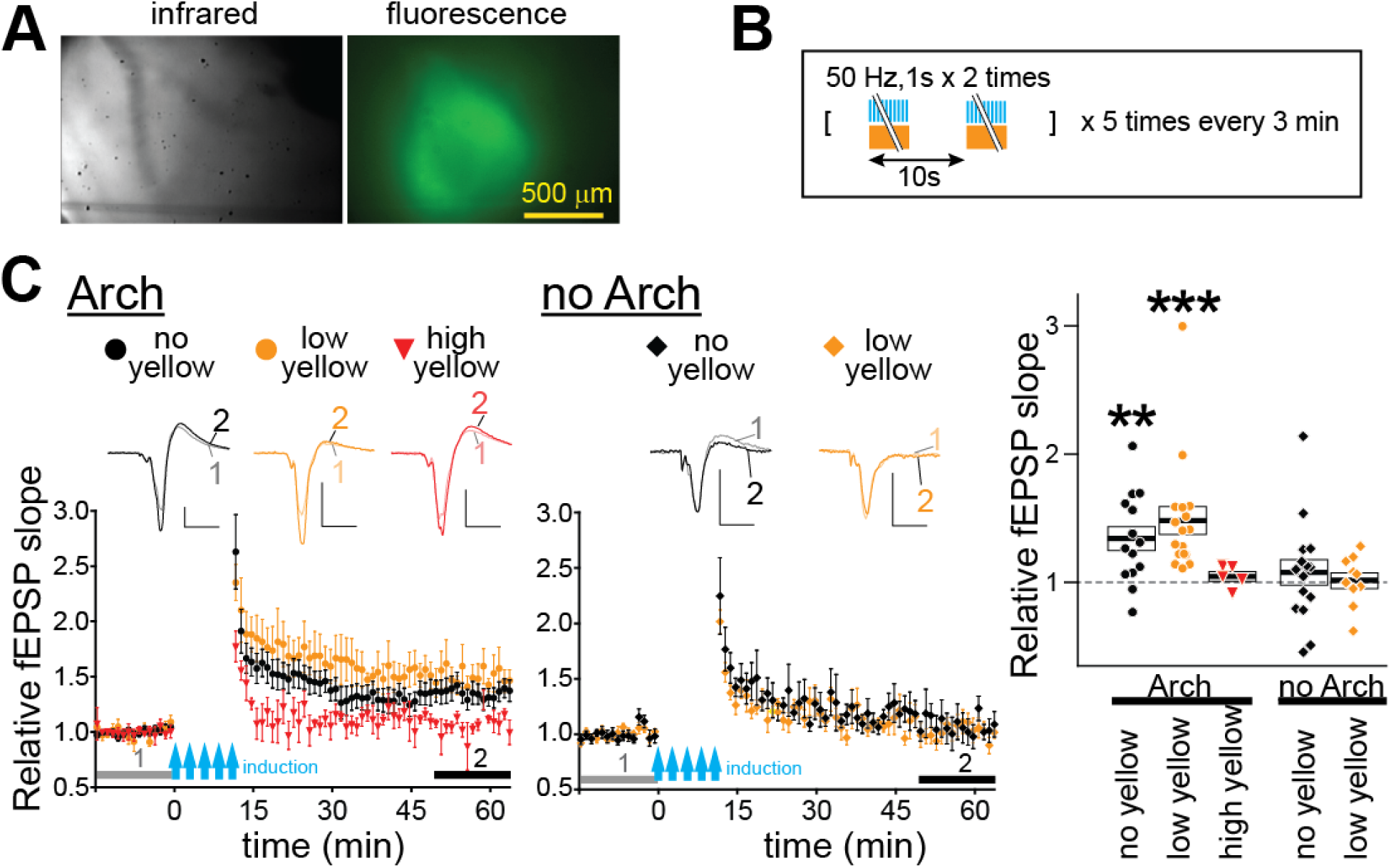
Arch suppression of Sst-INs enables LTP induction. (A) Right: Chronos-GFP and Arch-GFP fluorescence in BLA of an Sst-Cre driver mouse transduced with a Chronos-AAV in dmPFC and a floxed Arch AAV in BLA. Left: an IR image of the same slice. (B) LTP induction protocol. (C) LTP experiments on slices with Arch (left) and without Arch (middle) in Sst-INs, with LTP induced using pulses of blue light alone (black circle: no yellow) or combined with the continuous yellow light of two intensities: 0.15 mW/cm^2^ (orange circles: low yellow) and 0.24 mW/cm^2^ (red inverted triangles: high yellow). Insets represent examples of averaged fEPSPs before (1) and after (2) LTP induction as indicated by horizontal grey and black bars. Scales: 0.2 mV, 10 ms. Right: Summary data for relative fEPSP slopes averaged during (2) for each slice. n=14 (Arch-no yellow), 17 (Arch-low yellow), 5 (Arch-high yellow), 16 (no Arch-no yellow) and 10 (no Arch-low yellow). **p<0.01, ***p<0.001, compared to baseline, Wilcoxon Signed Rank Test. Boxes and the thick bars inside represent SEM and means.

LTP induction in the dmPFC-BLA input was tested by giving trains of blue light pulses alone (Fig.3A) or combined with the continuous yellow light of different intensities (Fig.4B). The yellow light by itself did not cause the release of glutamate from the dmPFC axonal terminals in BLA (data not shown). Unexpectedly, the trains of blue light in the absence of yellow light induced a significant LTP (Fig.4C, black-filled circles). Combining the trains of blue light and the yellow light of low intensity (0.15 mW/mm^2^) produced more significant LTP and with a tendency to be higher than with the blue light alone (Fig.4C, orange-filled circles). Increasing the yellow irradiance to 0.24 mW/mm^2^ prevented LTP induction (Fig.4C, red-filled inverted triangles). In slices without Arch, the low-intensity yellow light did not enable LTP induction by the pulses of blue light (Fig.4C, middle). Together, these data indicate that Arch enables LTP induction by the trains of blue light and it occurs even in the absence of yellow light, suggesting that the blue light inhibits Sst-INs expressing Arch. Consistently, whole-cell recordings from an Sst-IN with Arch revealed hyperpolarizing currents elicited by blue light (Fig.S2).

To test LTP induction *in vivo*, mice expressing Arch in the BLA Sst-INs and Chronos in dmPFC were implanted with optrodes, whose two electrodes were positioned in BLA and an optical fiber above BLA (Fig.5AB). The LTP induction protocol, identical to the protocol used *ex vivo*, but repeated 3 times with the one-hour interval, produced LTP, which lasted for almost 10 h (Fig.5CD).

**Fig.5.**
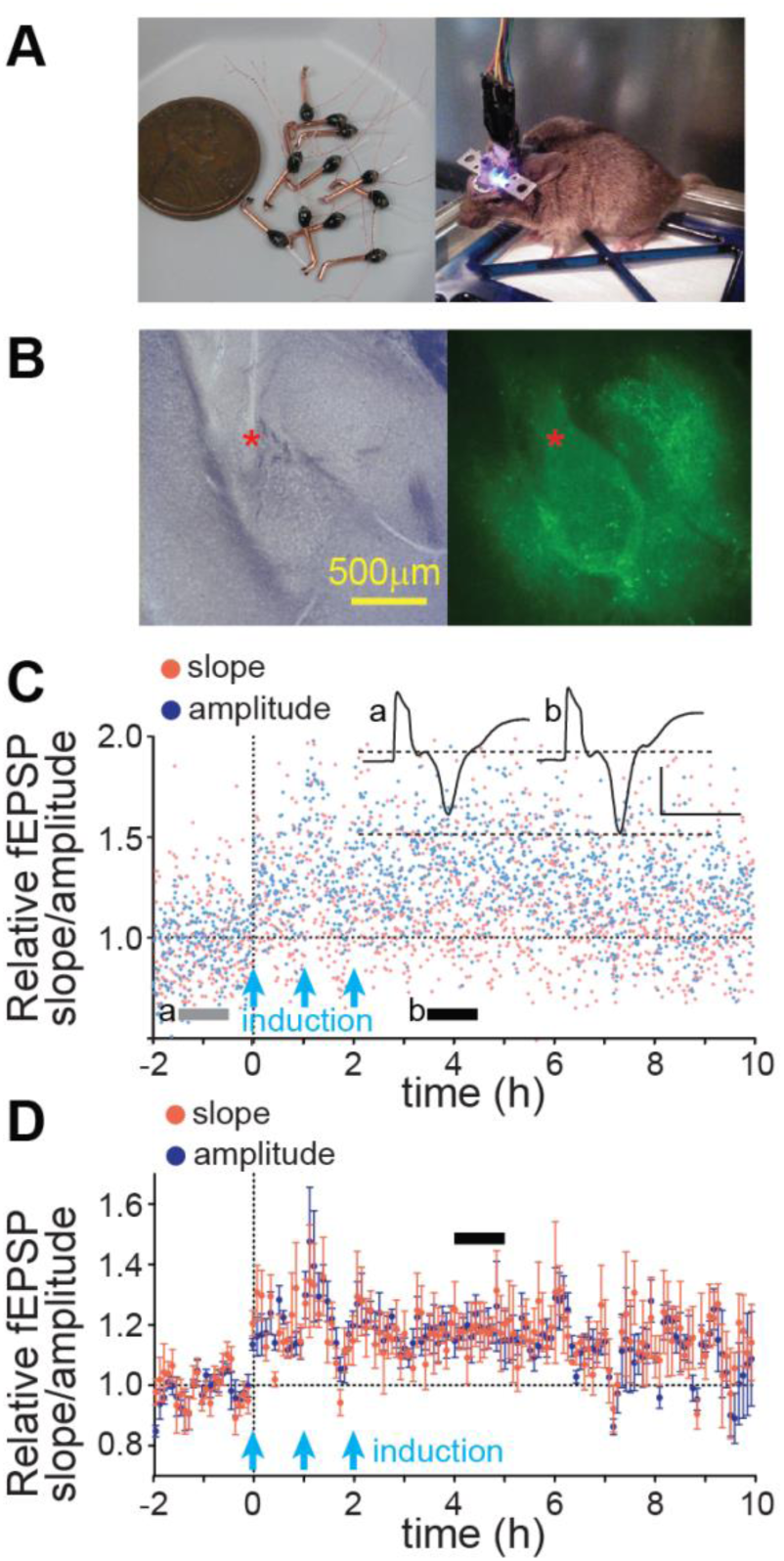
Arch-assisted LTP induction *in vivo*. (A) Left: LED light source-optrode assemblies, right: a mouse implanted with two optrodes aiming bilaterally at BLA. (B) An example position of electrodes (red asterisks) in a BLA slice imaged under the visible (left) or fluorescent (right) light. The fluorescence arises from Chronos-GFP in dmPFC axons and Arch-GFP in Sst-INs. (C) An example of an LTP experiment showing the slope (red) and amplitude (blue) of light-evoked fEPSP. Light-blue arrows show the trains of 50Hz light stimulation. The horizontal grey/black bars indicate the ranges for averaging for the sweeps shown in insets (a: grey, before LTP induction. b: black after induction). Scales: 0.4 mV, 5 ms. (D) Summary LTP data (n=6). The slope facilitation during the 3d hour after induction, identified by the horizontal bar, is highly significant (p<0.0001).

## 4 Discussion

This study has three findings: 1) Sst-INs, but not PV INs gate the plasticity in the dmPFC-BLA pathway induced by high-frequency presynaptic stimuli, 2) removal of the inhibition from Sst-INs, either chemogenetically or optogenetically, both enables the artificial facilitation of this pathway *ex vivo* and *in vivo*, and 3) blue light alone is sufficient for the optogenetically-assisted facilitation because the wavelength partially activates Arch expressed in Sst-INs.

The finding that suppression of Sst-INs, but not PV-INs, enables LTP induction in BLA input from dmPFC, suggests that Sst-INs are distinct groups of GABAergic neurons, specializing in gating synaptic plasticity in remote inputs to BLA. A similar role of Sst-INs was reported in the somatosensory cortex, where Sst-INs gate LTP in the lemniscal sensory pathway and their suppression by PV- and VIP-INs “opens that gate” and allows LTP induction (30). The LTP gating by Sst-INs, however, is not a universal phenomenon throughout the brain. For example, in the hippocampus, the Sst-INs located in the oriens/alveus region of the area CA1 rather enhance LTP in the Schaffer collateral pathway by inhibiting GABAergic neurons in the stratum radiatum and thereby disinhibiting the CA1 principal cells (31). The PV-INs, in turn, appear to gate the hippocampal LTP, based on the finding of a stronger LTP in the model mice for the pre-symptomatic ASL and Altzheimer, in which a mutated NRG1 receptor Erb4 causes deficiencies of the PV-INs (32-34). Thus, the PV-INs and Sst-INs oppose each other in both BLA and hippocampus, yet the roles of each IN population in LTP are reversed between the structures. Given such region-dependency of the PV- and Sst-IN functional relationship, the disinhibition-assisted LTP in different target regions may require suppressing of different subclasses of GABAergic neurons.

Another technical aspect is the choice between chemogenetic and optogenetic suppression. While DREADD is highly effective in suppressing GABA release from INs even when they fire action potentials, the drawbacks for *in vivo* experiments are the long washout times and the off target-effects of chemogenetic ligands (35). The optogenetic suppression of INs avoids these problems but using two different wavelengths of light *in vivo* is more expensive and technically demanding. An additional drawback of the two-light design is that stronger yellow light interferes with LTP induction. It suggests that yellow light desensitizes Chronos even at the levels that do not activate Chronos to release neurotransmitter, as seen with the ReaChR (36). Fortunately, this problem can be avoided by using blue light alone, which is sufficient for LTP induction both *ex vivo* and *in vivo* with Arch expressed in Sst-INs. This is because the blue light (470nm) still activates Arch, even though at the 35% efficiency of the yellow light (560nm) of the same power (37). Perhaps, the single-color disinhibition assisted LTP can be improved further by replacing Arch with a blue-shifted inhibitory opsin, like a proton pump Mac, activated by the blue 470 nm light at about 60% of the maximum efficiency (37).

## 6 Acknowledgments

The study was supported by NIH grants MH118604 and MH120290.

