## Supplemental Figures 1 and 2 for "Disinhibition-assisted LTP in the prefrontal-amygdala pathway via suppression of somatostatin-expressing interneurons"

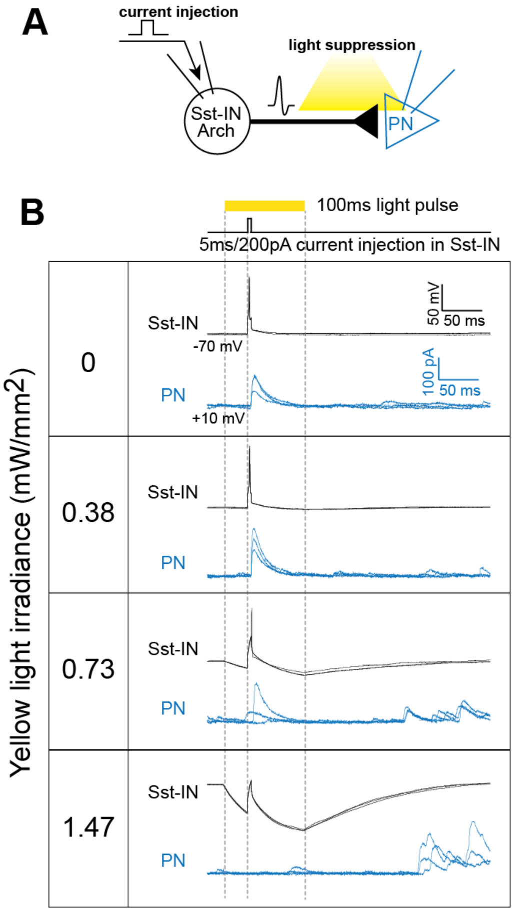


**Fig.S1 Arch inhibits GABAergic output from an Sst-IN.** (A) Experimental scheme. The 5 ms 200 pA depolarizing current was injected in an Sst-IN (at -70 mV) expressing Arch during the 100 ms pulse of yellow light, which started 30 ms before the current injection. Inhibitory postsynaptic currents were recorded from a connected principal neuron held at +10 mV. (B) Membrane potentials in the Sst-INs (black) and postsynaptic currents in the principal neuron (PN, blue) at different levels of yellow light irradiance.

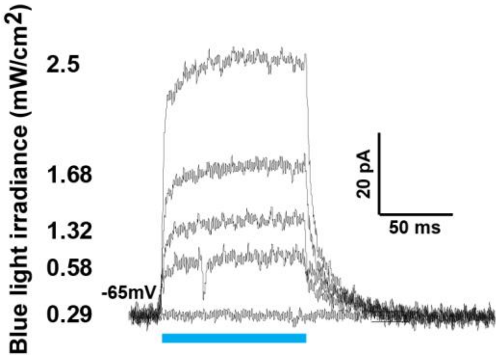


**Fig.S2 Blue light elicits hyperpolarizing current in an Sst-IN expressing Arch.** An example of the whole-cell recording from a BLA Sst-INs with Arch. The responses to 100 ms pulses of blues light of the indicated irradiance are shown.
